# Embodied Singing: Dual Role of Interoception in Vocal Expertise and Musical Competence

**DOI:** 10.1101/2025.04.14.648736

**Authors:** A.M. Zamorano, N. T. Haumann, D. Ligato, P. E. Maublanc, E. Brattico, P. Vuust, B. Kleber

## Abstract

Musical expertise is often associated with heightened perceptual sensitivity to external sensory stimuli, yet its relationship with internal bodily awareness (interoception) remains elusive. This study examined whether interoceptive ability relates differentially to varying levels of singing expertise and explored if interoception could predispose individuals to musical skills. Professional singers, amateur singers, and non-singers completed a heartbeat discrimination task (interoceptive accuracy; IAcc), self-reported interoceptive sensibility assessment, and comprehensive musical competence measures. Results demonstrated a significant positive association between singing expertise and IAcc, which notably emerged only in professional singers, who significantly outperformed both amateur singers and non-singers. Regression analyses indicated a moderate predictive role of accumulated singing practice for IAcc among trained singers, though emotional awareness partially mediated the relationship between expertise and IAcc. Critically, higher IAcc in non-singers significantly correlated with superior singing accuracy, suggesting enhanced interoception may facilitate singing competence independently of formal vocal training. Altogether, these findings highlight a dual role of interoception, linking it to expert-level singing—partially mediated by emotional awareness— and independently to musical competence in non-singers. These results reconcile prior inconsistencies by highlighting the embodied nature of singing and the distinct role of musical expertise, while underscoring inherent musically relevant mechanisms beyond practice alone.

## 1. Introduction

Interoception refers to the brain’s representation of internal bodily activity, such as muscle tension, heart rate, respiratory effort, temperature, and pain^1,2^. These signals are processed in brain regions involved in predicting and representing information from the body, including the brainstem, insula, anterior cingulate cortex and ventromedial prefrontal cortex^2,3^. Interoception is crucial for a range of physiological and cognitive functions, such as autonomic regulation, sensorimotor regulation, higher-order cognition, and emotion processing ^4^. Consequently, a growing body of research has demonstrated a link between the perception of internal bodily signals and affective states, such as anger, joy, aversion, or hunger^5,6^. Yet, despite evidence linking interoception to affective and cognitive processes, it remains unclear how heightened interoceptive ability relates to skilled motor behaviors—such as singing—where precise sensory-motor integration and bodily self-monitoring are crucial.

Musicians’ extensive and disciplined training, coupled with precise integration of sensory and motor processes, is an exemplary model for investigating the brain’s abilities to adapt to movement-related and perceptual skill development^7–9^. Among musicians, singers present a unique opportunity to understand the interplay between body and brain, as they rely on the intimate connection between their physiology and sound production^10–12^. Specifically, singing requires volitional breath control, precise laryngeal muscle coordination to regulate vocal fold tension, and vocal tract shaping to achieve stylistically appropriate pitch, timbre, and dynamics. The continuous physiological awareness of airflow, muscle engagement, and articulator shape enables singers to fine-tune these mechanisms to optimize acoustic output and ensure controlled, expressive vocal performance^13–15^. Such heightened reliance on internal bodily signals makes singing a deeply embodied activity, requiring the sophisticated integration of bodily inputs, higher-order cognition, and motor control to anticipate musical structures and regulate performance dynamics with precision^16–18^.

Empirical findings suggested that extensive dedicated musical training can enhance interoceptive ability, indicating that musicians with more extensive training may have greater interoceptive accuracy (IAcc) compared to non-musicians^19,20^. However, conflicting evidence in a recent cross-sectional and longitudinal study testing the role of musical training in increasing interoception challenged that notion^21^. This lack of replication raises questions about the extent and nature of enhanced interoceptive perception in musically trained individuals. Specifically, it remains unclear whether interoception is shaped by the level of musical expertise cultivated through extensive training (nurture) or if it is an innate ability (nature) that predisposes individuals to excel in musical performance. Furthermore, it is uncertain whether enhanced interoceptive accuracy plays a role in musical abilities or whether it merely correlates with varying levels of expertise and years of dedicated practice.

This study aims to explore these questions by examining the relationship between IAcc and musical expertise across varying levels of singing proficiency. To that end, we compare non-singers, amateur singers, and professional singers, assessing their heartbeat discrimination ability alongside their performance on musical competence tasks and self-reported interoceptive sensibility. The heartbeat discrimination task is a well-established, objective physiological measure for evaluating individual differences in IAcc^22^, whereas interoceptive sensibility measures one’s subjective perception and regulation of bodily signals across multiple dimensions^23^. We hypothesize that notable differences in IAcc arise only in individuals with advanced musical training. Specifically, we expect professional singers—who undergo extensive vocal training and sensorimotor refinement—to show greater IAcc than amateur singers and non-singers. Conversely, if no significant group differences are observed, this may indicate that IAcc is a relatively stable trait that is not strongly influenced by musical training. Finally, if IAcc correlates with musical competence tasks in non-singers, this would suggest that interoceptive ability may act as a predisposing factor for musical skill development, highlighting a potential interaction between training effects and pre-existing individual differences.

## 2. Materials & Methods

### 2.1 Participants

Participants were recruited online and through advertisements at Aarhus University and The Royal Academy of Music, Aarhus/Aalborg. Initially, 74 participants were recruited across three groups: professional singers, amateur singers, and non-singers. Final sample details, including any exclusions, are reported in the Results section (see Table 1). Exclusion criteria included any history of neurological, cardiorespiratory, or mental disorders, as well as pregnancy. For non-singers, additional exclusion criteria included any prior singing training or regular singing engagement (e.g., choir activities).

**Table 1:**
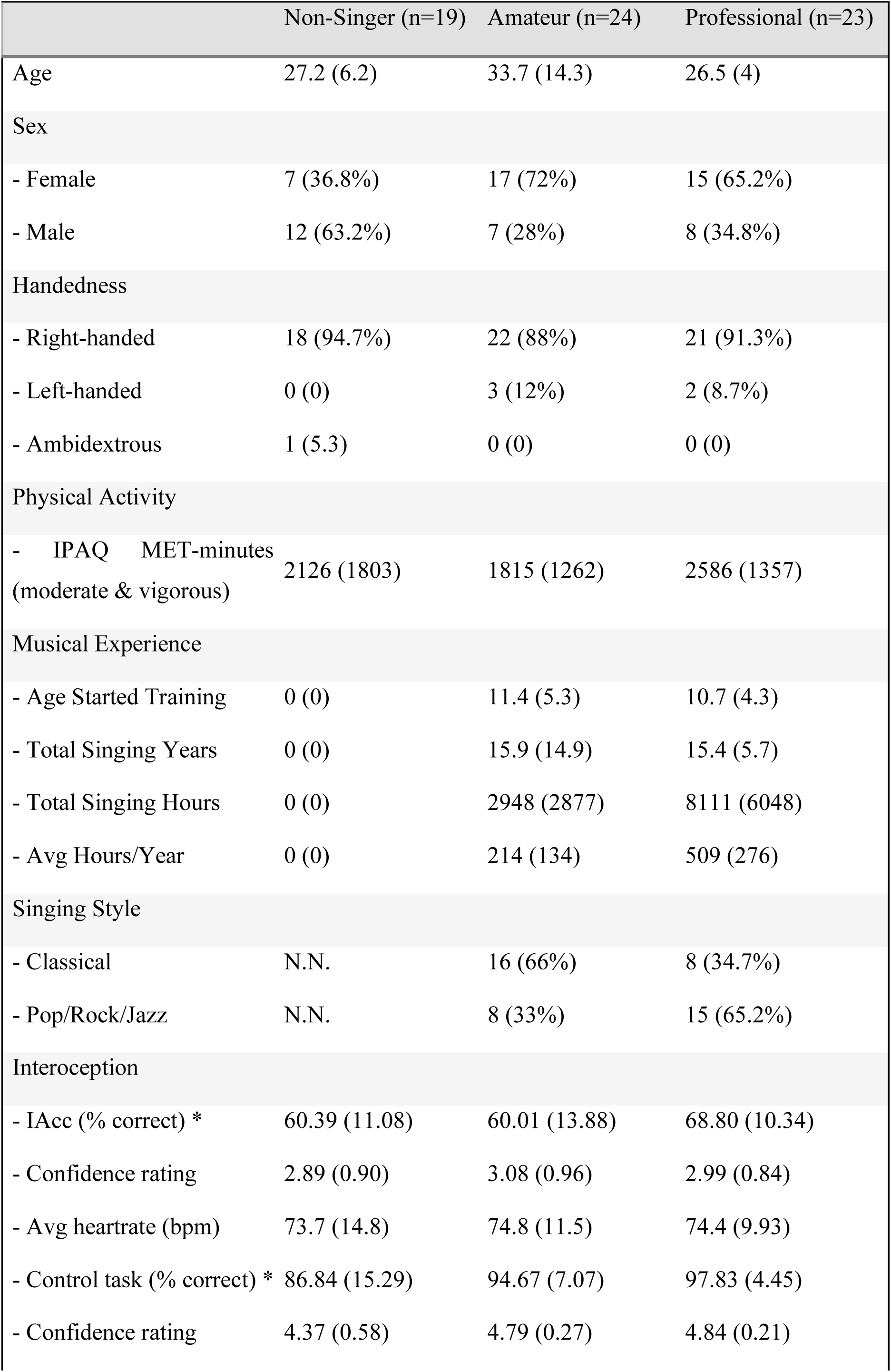
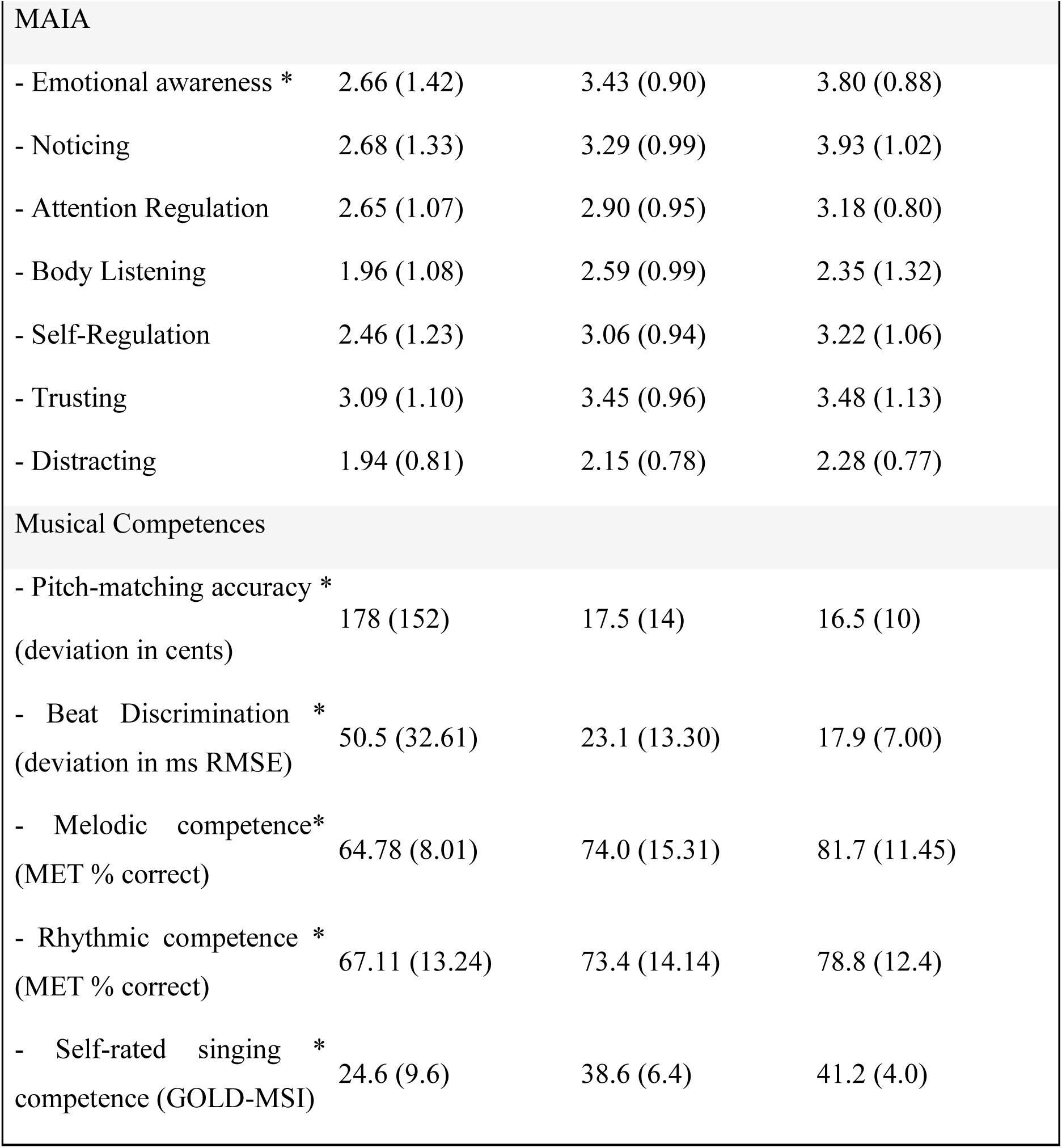
Demographic characteristics, musical background, interoceptive measures, and musical competence across non-singers, amateur singers, and professional singers. Values represent means (standard deviations) or frequencies (percentages). Interoceptive accuracy (IAcc) reflects performance (% correct responses) on the heartbeat discrimination task; interoceptive sensibility was assessed using the Multidimensional Assessment of Interoceptive Awareness (MAIA). Musical competence was assessed using a Singing Accuracy task (Pitch-Matching Accuracy, measured as deviation from target pitch in Cent), equidistant beat discrimination accuracy (RMSE; ms), the Musical Ear Test (MET, melodic [METm] and rhythmic [METr] subtests), and self-rated singing competence (Goldsmiths Musical Sophistication Index, GOLD-MSI Singing Competence subscale). Physical activity levels were measured using the International Physical Activity Questionnaire Short Form (IPAQ-SF; moderate and vigorous MET-minutes per week). Significant group differences (p < 0.05) are indicated by asterisks (*).

Sample size determination was conducted using GPower 3.1^24^, based on previous studies investigating interoceptive accuracy in performing artists^19,21,25^, targeting to power the study at 80% to detect an effect size of 0.40 in a one-way ANOVA (α = 0.05). All participants were verbally informed about the study details and provided written consent before participation. The study was conducted in accordance with the Declaration of Helsinki and approved by the Scientific Ethics Committee for Region Midtjylland (Komité II; #1-10-72-65-17).

### 2.2 Experimental design

Participants attended a single session to assess interoception, musical competence, and self-reported traits. They sat comfortably with legs uncrossed and arms resting without bodily contact. Interoceptive accuracy and a perceptual control task were prioritized to prevent potential cognitive fatigue from affecting interoception and confidence ratings, while music competence tasks were performed later, avoiding any influence of IAcc performance through priming effects. The order in which the interoception and control tasks, as well as the respective music competence tasks were presented was counterbalanced. Self-report data were collected online using SurveyXact (Ramboll University, Denmark) once all the tasks were completed.

#### 2.2.1 Interoception

*Interoceptive Accuracy* was assessed using a heartbeat discrimination paradigm^22,26^, where participants judged whether external auditory tones were presented synchronously or asynchronously with their heartbeat. The paradigm consisted of 30 trials, each delivering 10 auditory tones (800 Hz, 100 ms duration). Tones were triggered by the participant’s ECG R-wave and presented at 250 ms (synchronous condition) or 550 ms (asynchronous condition) post-R-wave for distinguishing heartbeat perception^27,28^. Participants simultaneously attended to their heartbeat and the tones, responding to: "Were the tones delivered synchronously or asynchronously with your heartbeat?" After each trial, they rated their confidence on a five-point Likert scale (1 = Not at all sure, 5 = Completely sure) in response to: "How confident are you in your response?", measuring metacognitive awareness of IAcc.

Electrocardiographic (ECG) signals were recorded (BrainAmp ExG, BrainProducts, Germany) via lead II chest electrode placement. The ECG signal was transmitted to a second laptop running MATLAB® and Brainvision RDA Client. Online R-peak detection used custom MATLAB code (25 Hz bandpass filter, automated threshold calibrated over a 10-second baseline). Detected R-peaks triggered audio signals sent via minijack to a third laptop running Presentation® software, with a processing delay of 203 ms (SD = 14) accounted for to ensure precise R-peak-to-tone timing.

*The Exteroceptive Control Task* required participants to judge whether external auditory tones were presented in or out of synchrony with a tactile stimulus, assessing whether group differences in IAcc might stem from temporal estimation biases. The task mirrored the IAcc task design, with participants responding to the same questions, evaluating tone synchrony with the tactile stimulus, and rating their confidence on a five-point Likert scale.

Tactile stimuli were delivered via a current generator (DeMeTec, Langöns, Germany) and a concentric surface electrode (WASP, Brainbox Ltd., UK) placed on the index fingertip. Tactile detection thresholds were established using a staircase procedure^29^ starting at 1mA, doubling until detection, then decreasing intensity until the sensation disappeared. The step-factor was set at 1.25, continuing until six reversal points were obtained. The geometric mean of the last eight reversals from two runs defined the final threshold. To ensure reliable stimulus detection, the task intensity was set 0.4mA above this threshold.

*Interoceptive Sensibility* was assessed using the Multidimensional Assessment of Interoceptive Awareness (MAIA) scale^30^. This 32-item self-report instrument evaluates various facets of interoception, including the ability to regulate attention and emotional responses to bodily signals, across eight dimensions: noticing, not-distracting, not-worrying, attention regulation, emotional awareness, self-regulation, body listening, and trusting. Notably, MAIA scores do not reliably predict objective interoceptive accuracy^23^, reflecting its emphasis on broader experiential aspects of interoception.

#### 2.2.2 Musical Competence

Music-related experience was assessed with the Montreal Music History questionnaire ^31^. The musical competence tasks aimed to clarify the relationship between interoception and musical abilities, assessing whether IAcc is functionally relevant to musical tasks or a byproduct of extensive sensorimotor training.

A *Vocal Pitch-Matching Task* was employed to objectively measure singing accuracy^12,16^, defined as pitch-matching accuracy, quantified as deviation (in cents) from target pitches. During this task, participants heard 54 pseudorandomized musical intervals presented via headphones using Max/MSP software (Cycling 74, San Francisco, USA). For each interval, participants reproduced two pitches vocally. The first tone was fixed at 311.13 Hz (D#4) for females and 155.565 Hz (D#3) for males, while the second tone was either identical (4×) or differed by 1 (6×), 3 (12×), 5 (12×), 6 (12×), or 7 (8×) semitones, equally distributed between ascending and descending intervals. Vocal responses were recorded and saved in wav format for offline automated analyses of pitch-matching accuracy. Pitch-matching accuracy was defined as the deviation between sung and target tones in cent (100 cent = 1 semitone).

*Equidistant Beat Discrimination Accuracy* was assessed with a custom-built experiment in JavaScript, designed to assess temporal estimation accuracy in the context of musical beats. In this task, two isochronous kick drum (KD) sounds were presented with a fixed inter-onset interval (IOI) of 1000 ms. A snare drum (SD) sound was presented between the two KD sounds, with its temporal onset adjustable relative to the KD sounds. Participants’ task was to determine whether the SD sound was "in beat"—defined as perfectly equidistant between the two KD sounds (500 ms)—or "out of beat." At the start of each block, participants were given an example of an equidistant beat. During trials, the SD sound was initially presented with a 650-ms offset to ensure clear out-of-beat perception. Participants adjusted the SD timing by clicking buttons indicating if it sounded "in beat" or "out of beat," following a one-up, two-down staircase procedure. Initial step size (25 ms) decreased by a factor of 1.25 after two consecutive positive responses, and further reduced by 1.15 after the second reversal. The task ended after 10 reversals, and perceptual thresholds were calculated as the root mean squared error (RMSE) over the last eight reversals.

*Music Perception* was assessed using the Musical Ear Test (MET)^32^, which comprises melodic (METm) and rhythmic (METr) tasks, each consisting of 52 trials. METm evaluates participants’ ability to discriminate between pairs of melodic phrases by presenting two short note sequences that are either identical or differ by one note. Participants determine whether the melodies are the same or different, emphasizing melodic discrimination and memory. METr assesses rhythmic pattern recognition, where participants compare two sequences of short and long note durations and judge whether they match, testing sensitivity to rhythmic variations and patterns.

*Self-perceived singing proficiency* was assessed using the Goldsmiths Musical Sophistication Index (Gold-MSI) Singing Competence subscale (Mullensiefen et al., 2014). This seven-item subscale measures agreement with statements about singing skills, including pitch accuracy and confidence in performance, on a 7-point Likert scale. It contributes to the overall musical competence profile by capturing aspects of vocal performance and self-assessed singing competence.

*Acoustic Frequency-Discrimination Thresholds* were measured to ensure that each participant possessed intact basic pitch perception skills, ruling out perceptual differences as a factor in production performance. These thresholds were assessed using a two-interval forced-choice paradigm^17^. In each trial, participants heard two 250-ms pure tones (10-ms ramps) separated by a 600-ms gap. A standard tone (500 Hz) and a target tone (500 Hz plus a frequency difference, fΔ) were presented in random order. Participants indicated which interval contained the higher-frequency target. A two-down one-up adaptive procedure estimated the 70.7% correct threshold on the psychometric function^33^. Each block began with a large fΔ (7%) for detectability. After two consecutive correct responses, fΔ decreased; after one incorrect response, it increased. The step factor, initially 2, was reduced to 1.25 after the second reversal. The procedure continued for 15 reversals, with the geometric mean of the final eight defining the threshold. Each participant completed two runs, and their average threshold was recorded as a percentage of the standard frequency.

### 2.3 Physical activity and affective traits

*Physical Activity* levels account for potential confounding effects on interoceptive accuracy, as prior research suggests a link between higher activity levels and enhanced IAcc^34,35^. Activity levels were assessed using the International Physical Activity Questionnaire - Short Form (IPAQ-SF)^36^, a validated instrument measuring physical activity over the past seven days. The IPAQ-SF includes nine items assessing time spent on vigorous activities (e.g., aerobics), moderate activities (e.g., leisure cycling), walking, and sitting. To control for extreme values, participants with excessive physical activity levels—defined as combined moderate and vigorous MET-minutes exceeding three standard deviations from the mean—were excluded post-hoc (see Results section).

*Affective Traits.* IAcc has been linked to affective processes, shaping the misinterpretation of physical symptoms as anxiety-related signals^37,38^. Given that performing artists are particularly susceptible to music performance anxiety (MPA)—which shares features with social anxiety^39^—we examined the relationship between affective traits and IAcc. General anxiety sensitivity was measured using the State-Trait Anxiety Inventory-Trait version (STAI-T)^40^, while social anxiety was assessed with the Mini Social Phobia Inventory (Mini-SPIN), a validated screening tool^41^.

### 2.4 Statistical analysis

All statistical analyses were conducted using JASP (version 0.19.2). Assumptions of normality and homogeneity of variance were assessed using Q-Q plots and Levene’s test, respectively. Where violations were detected, the corresponding robust post-hoc tests and adjustments to degrees of freedom were applied, and the appropriate analysis are reported in the results section. Effect sizes, including partial eta squared (η²), Cohen’s d, and Dunn’s Rank-Biserial Correlation are reported to facilitate interpretation. Statistical significance was set at p < 0.05. Participants with incomplete or invalid responses and those meeting predefined exclusion criteria were excluded from the analyses. Outliers were identified using z-scores exceeding ±3 SD to ensure the robustness of statistical conclusions.

#### Group Comparisons

Group differences in IAcc, confidence ratings, the exteroceptive control task, Multidimensional Assessment of Interoceptive Awareness (MAIA), self-reported physical activity (IPAQ), and affective traits (e.g., STAI-T and Mini-SPIN) were assessed using ANCOVAs, with singing experience (non-singer, amateur, professional) as the independent variable. Age and sex were included as covariates to account for potential confounding effects. Post-hoc pairwise comparisons were conducted using Tukey’s HSD correction for equal variances and Games-Howell correction for unequal variances, ensuring appropriate control of family-wise error rate while accounting for variance homogeneity assumptions. A multivariate analysis of covariance (MANOVA) was conducted to examine group differences in musical competence outcomes. Following a significant MANOVA, univariate ANOVAs with Tukey’s HSD post-hoc tests identified specific group differences.

#### Relationships Between Variables

To examine the relationship between IAcc, singing experience, and music competence, and affective traits, multiple approaches were employed, including Regression Analysis, Partial Correlations and Mediation Analyses. Mediation Analyses were performed using bootstrapping with 5,000 resamples to estimate direct, indirect, and total effects. Path analyses were employed to quantify mediation effects, with direct effects representing the relationship between variables after accounting for mediators, and indirect effects capturing the proportion of variance explained by the mediators.

## 3. Results

### 3.1 Participants

The final sample comprised 67 participants (27 males, 40 females) distributed across non-singers (n = 19, 12 males, 7 females), amateur singers (n = 24, 7 males, 18 females), and professional singers (n = 23, 8 males, 15 females).

Four participants were excluded from statistical analyses due to compromised data quality: Two non-singers did not adhere to task instructions, with one terminating the experiment prematurely and the other providing invalid responses by repeatedly pressing the same key across trials. Two professional singers were excluded, as one reported mild discomfort the day of the experiment, and the other reported difficulty focusing on the tasks after insufficient rest following a late-night performance.

Outliers were identified using conservative z-score thresholds exceeding ±3 SD, leading to the exclusion of another four participants: A female non-singer was identified with excessive daily high-level endurance exercise, our primary exclusion criterion based on its influence on interoceptive accuracy^34,35^. Additionally, one female non-singer with an abnormally low IAcc score of only 7% correct responses and a male amateur singer with insufficient pitch accuracy for this group (> 2 semitones) were removed from the analyses.

A detailed description of the final sample is provided in Table 1.

### 3.2 Interoceptive accuracy

An ANCOVA was conducted to examine the effect of singing experience on cardiac interoceptive accuracy (IAcc), while controlling for age and gender. Assumption checks indicated no significant violation of homogeneity of variances, as assessed by Levene’s test, *F*_(2, 63)_ = 0.68, *p* = 0.51, and the Q-Q plot suggested that the residuals were approximately normally distributed. The ANCOVA revealed a significant main effect of singing experience on interoceptive accuracy, *F*_(2, 61)_ = 4.18, *p* = 0.02, with a partial eta squared (*η²*= 0.12), suggesting a moderate effect. Neither age (*F*_(1, 61)_ = 0.19, *p* = 0.66) nor sex (*F*_(1, 61)_ = 1.07, *p* = 0.30) were significant covariates. Post-hoc analyses confirmed that professional singers exhibited higher IAcc than non-singers (*p* = 0.04, *d* = 0.78) and amateurs (*p* = 0.04, *d* = 0.76) with moderate to large effect sizes (Figure 1A). The difference between amateurs and non-singers was not significant (*p* = 1.00, *d* = −0.02).

**Figure 1:**
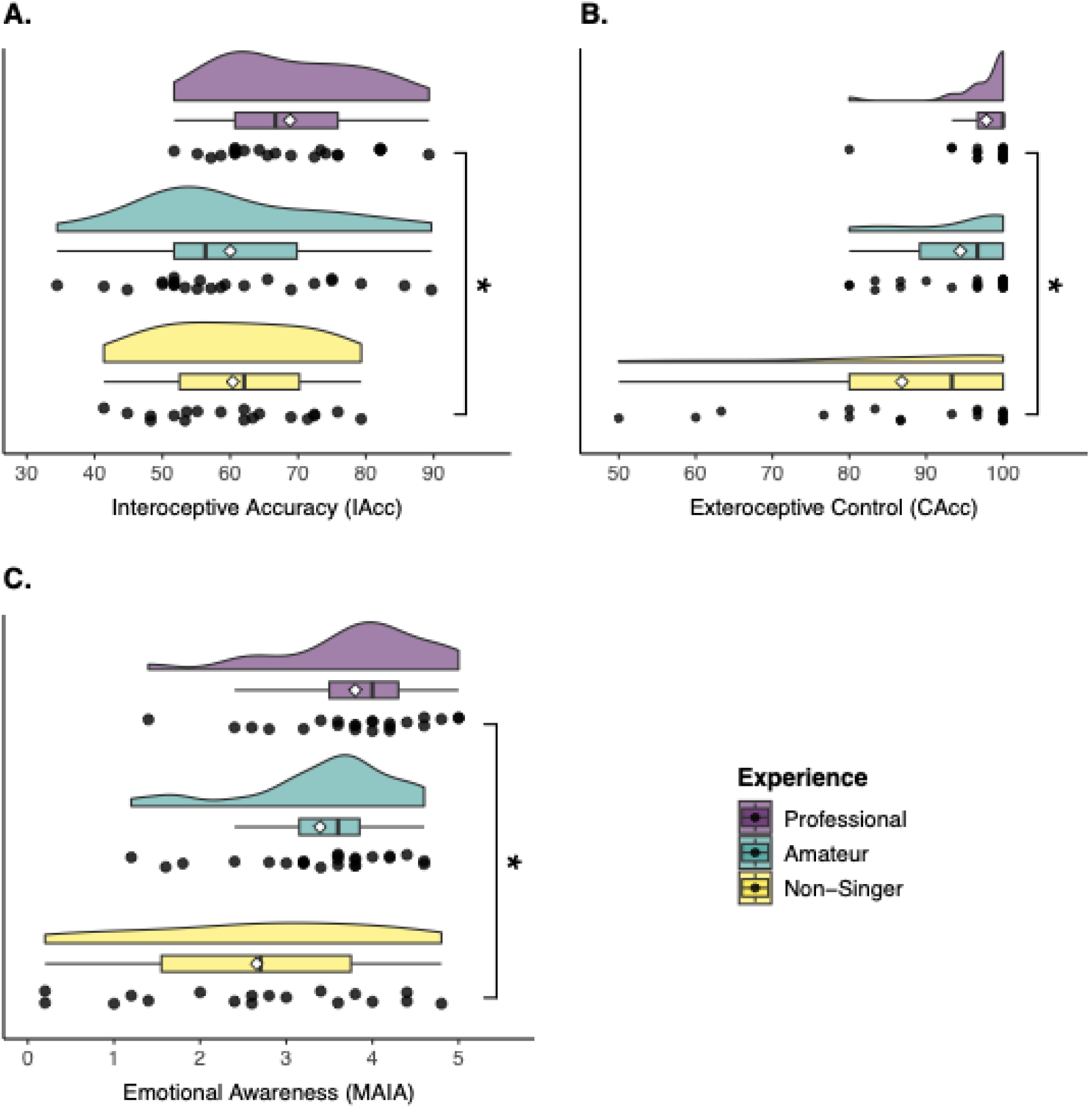
Raincloud plots illustrating (A) Interoceptive Accuracy (IAcc), (B) Exteroceptive Control Task performance, and (C) Emotional Awareness (MAIA subscale) across groups with varying levels of singing experience (non-singers, amateurs, professionals). Each plot displays the distribution of individual participant scores, median, interquartile range, and group means (depicted as diamond shapes). (A) Professionals exhibited significantly higher IAcc compared to non-singers, while no significant difference was observed between amateurs and non-singers. (B) The Exteroceptive Control Task required participants to judge whether external auditory tones were presented in or out of synchrony with a tactile stimulus, assessing temporal estimation biases. Professionals significantly outperformed non-singers, whereas amateurs did not differ significantly from either group. (C) Professionals reported significantly higher Emotional Awareness compared to non-singers, while no significant differences were observed between professionals and amateurs or between amateurs and non-singers. Significant between-group differences, identified through post-hoc tests, are marked with asterisks (*). Raincloud plots were generated using the ggrain package in R ^42^.

An ANCOVA was conducted to examine group differences in mean confidence ratings for interoceptive accuracy (IAcc), while controlling for age and sex. The analysis revealed no significant main effect of singing experience on confidence ratings, *F*_(2, 61)_ = 0.82, *p* = 0.44, *η²*= 0.03. Age was not a significant covariate (*F*_(1, 61)_ = 0.03, *p* = 0.87, *η²*= 0.00). However, sex showed a significant effect on confidence ratings, *F*_(1, 61)_ = 5.80, *p* = 0.02, *η²*= 0.09, suggesting that it accounted for some variance in confidence ratings irrespective of singing experience. A Pearson’s partial correlation examined the relationship between interoceptive accuracy (IAcc) and confidence ratings, while controlling for age and gender. This analysis revealed a positive but non-significant association, *r*= 0.23, *p* = 0.07.

### 3.3 Exteroceptive control task

To investigate the effect of singing experience (non-singers, amateurs, professionals) on temporal estimation judgements in the tactile-tone synchrony task, an ANCOVA was initially performed, indicating that neither age nor gender contributed significantly to the model. However, as Levene’s test revealed a violation of homogeneity of variances, *F*_(2, 63)_ = 14.15, *p* < 0.001, a non-parametric Kruskal-Wallis non-parametric test with Dunn’s post-hoc test (Bonferroni-corrected) was employed instead (Figure 1B). The analysis revealed a significant main effect of singing experience (H(2) = 8.46, p = 0.015, η² = 0.10). Dunn’s post hoc test indicated that professionals significantly outperformed non-singers (*p* = .011, r_rb = 0.47, large effect), while amateurs did not significantly differ from professionals (*p* = .454, r_rb = 0.24, medium effect) nor from non-singers (*p* = .348, r_rb = 0.15, small effect).

### 3.4 Interoceptive Sensibility

Partial correlations were conducted initially to explore the relationship between IAcc and Multidimensional Assessment of Interoceptive Awareness (MAIA) subscales, while controlling for singing experience, age, and sex. A significant positive correlation emerged between IAcc and the MAIA Emotional Awareness subscale (*r* = 0.26, *p* = 0.04), indicating that individuals with greater emotional awareness also demonstrated higher interoceptive accuracy, regardless of singing expertise. No significant relationships were observed with other MAIA subscales, including Noticing (*r* = 0.18, *p* = 0.15), Attention Regulation (*r* = 0.11, *p* = 0.39), and Body Listening (*r* =0.08, *p* = 0.56), nor with Self-Regulation, Trusting, or Not Distracting (all *p* > 0.40).

A subsequent ANCOVA was initially conducted to further examine the effect of singing experience (non-singers, amateurs, professionals) on the MAIA Emotional Awareness subscale, indicating that neither age nor gender contributed significantly to the model. However, as Levene’s test revealed a violation of homogeneity of variances, F(2, 62) = 3.72, *p* = 0.03, a non-parametric Kruskal-Wallis non-parametric test with Dunn’s post-hoc test (Bonferroni-corrected) was employed instead (Figure 1C). The analysis revealed a significant main effect of singing experience, (H(2) = 6.11, p = 0.047, η² = 0.11). Dunn’s post hoc tests revealed significantly higher emotional awareness in professionals compared to non-singers (p = 0.011, rank-biserial correlation = 0.49, large effect). No significant differences were found between amateurs and professionals (p = 0.364, r_rb = 0.24) nor between non-singers and amateurs (p = 0.135, r_rb = 0.15).

### 3.5 Musical Competence

A multivariate analysis of variance (MANOVA) was conducted to examine group differences in Singing Accuracy (Pitch-Matching Accuracy, deviation from target pitch in Cent), Equidistant Beat Discrimination Accuracy (RMSE, ms), Melodic Competence (METm), Rhythmic Competence (METr), and self-reported singing ability (GOLD-MSI) across three singing experience groups (non-singers, amateurs, professionals). Pillai’s Trace was used due to its robustness against violations of homogeneity of covariance assumptions. The MANOVA revealed a significant multivariate effect of singing experience, Pillai’s Trace = 0.757, F(10, 114) = 6.944, *p* < .001.

Follow-up univariate ANOVAs confirmed significant effects of singing experience across all tasks (Table 1, Figure 2). Levene’s test indicated violations of homogeneity of variances for Pitch Matching Accuracy (*p* < .001), RMSE (*p* < .001), METm (*p* = .0005), and GOLD-MSI (*p* = .004), necessitating the use of Games-Howell corrections for post-hoc comparisons in these cases. For METr (*p* = .23), equal variances were assumed, and Tukey’s HSD was applied. Effect sizes are reported as η² for ANOVAs and Cohen’s d for post-hoc comparisons. The results were as follows:

- *Pitch Matching Accuracy (Figure 2A)*: *F*_(2,63)_=26.09, *p*<0.001, η² = .45. Professionals and amateurs deviated significantly less from the target pitch (in Cent) compared to non-singers (both *p* < 0.001, *d* = 1.96–1.97), with no difference between professionals and amateurs (*p* = 0.99, *d* = 0.02).
- *Equidistant Beat Discrimination Accuracy (RMSE; (Figure 2B)*: *F*_(2,60)_=15.62, *p* < 0.001, η² = .34. Professionals and amateurs outperformed non-singers (both *p* < 0.01, *d* = 1.43–1.70), with no significant difference between professionals and amateurs (*p* = 0.64, *d* = 0.27).
- *Melodic Competence (METm; Figure 2C)*: *F_(2,63)_* = 9.78, *p* < 0.001, η² = .24. Professionals scored higher than amateurs and non-singers (both *p* < 0.01, *d* = 1.37– 1.42), while amateurs outperformed non-singers (*p* = 0.06, *d* = 0.72).
- *Rhythmic Competence (METr; (Figure D)*: *F*_(2,63)_ = 4.10, *p* = 0.02, η² = .12. Professional singers outperformed non-singers (*p* = 0.02, *d* = 0.88). Amateurs did not differ significantly from either group (*p* > 0.05, *d* = 0.42–0.46).
- *Self-reported Singing Ability (GOLD-MSI; (Figure 2E)*: *F*_(2,62)_ = 33.04, *p* < 0.001, η² = .52. Professionals scored higher than amateurs and non-singers (both *p* < 0.001, *d* = 2.43–2.57), with amateurs also outperforming non-singers (*p* < 0.001, *d* = 1.43).

**Figure 2:**
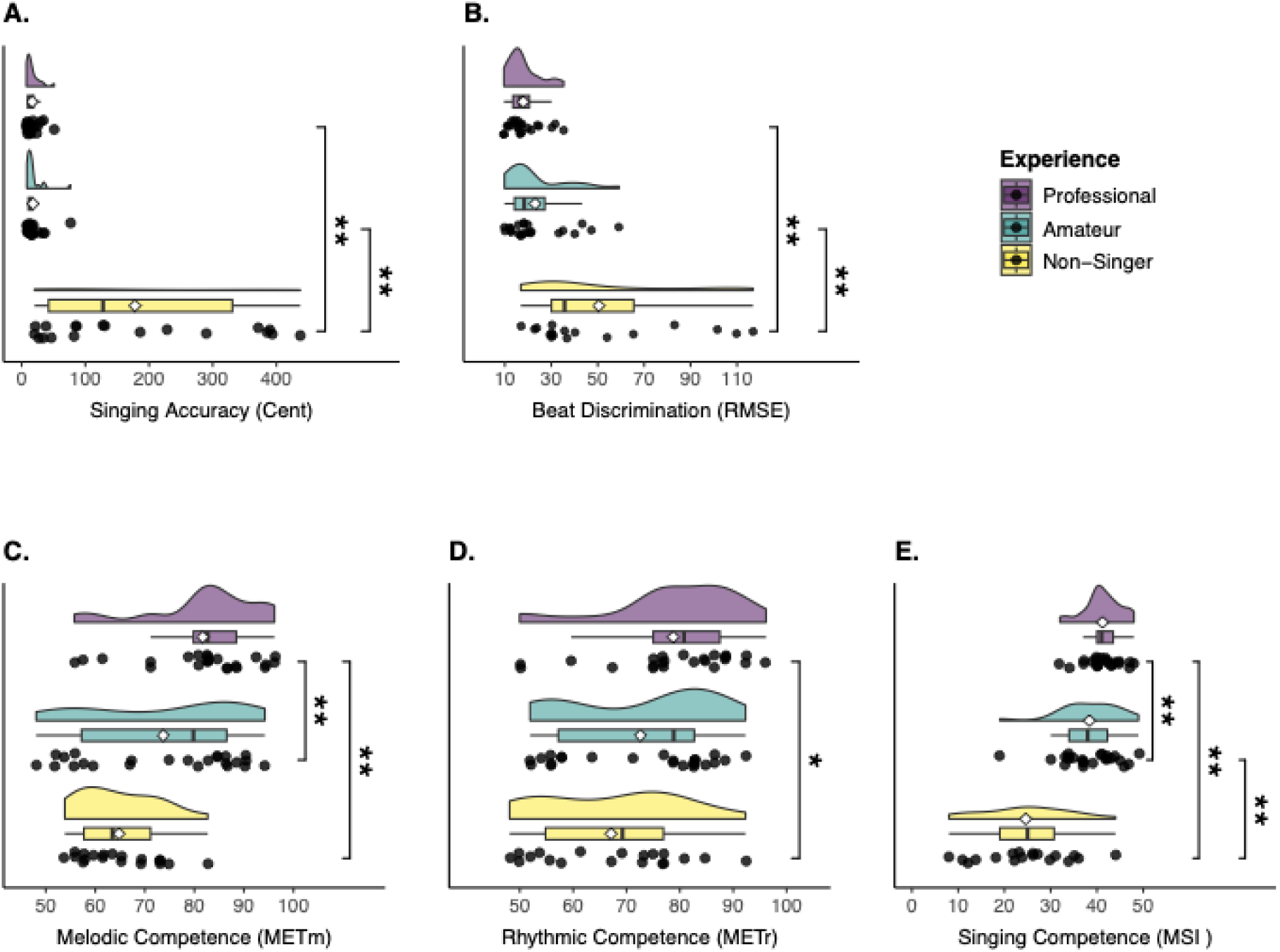
Raincloud plots illustrating group differences in musical engagement and performance across five tasks: (A) Pitch Matching Accuracy (deviation from target pitch in Cent), (B) Equidistant Beat Discrimination Accuracy (RMSE; ms), (C) Melodic Competence (METm), (D) Rhythmic Competence (METr), and (E) Self-reported Singing Competence (GOLD-MSI Singing Scale). Each plot displays data for non-singers, amateurs, and professional singers, showing the distribution of individual participant scores, median, interquartile range, and group means (depicted as diamond shapes). Lower scores indicate better performance for Pitch Accuracy and Beat Discrimination Accuracy, with RMSE representing the mean squared error of beat alignment in milliseconds (ms). In contrast, higher scores indicate better performance for METm, METr, and the GOLD-MSI Singing Scale. Significant between-group differences, identified through post-hoc tests, are marked with asterisks (*). Raincloud plots were generated using the ggrain package in R ^42^.

To ensure that differences in pitch perception could not account for the observed group differences in singing accuracy, a frequency discrimination task was included as a control measure from the outset. This task was designed to assess whether variations in singing accuracy were attributable to differences in basic frequency perception rather than vocal-motor control. The average frequency discrimination threshold (geometric mean) was 0.57% (min = 0.12%, max = 0.99%, approximately 9.88 cents) in professional singers, 0.51% (min = 0.08%, max = 0.93%, approximately 8.77 cents) in amateur singers, and 1.22% (min = 0.39%, max = 3.33%, approximately 20.67 cents) in non-singers. These results confirm that all participants demonstrated at least good to average pitch perception accuracy, ruling out the possibility that perceptual deficits contributed to the observed differences in pitch-matching performance. This interpretation is further supported by a non-significant correlation between pitch discrimination ability and deviations from target pitch for pitch-matching in non-singers (*r* = -0.11, *p* = 0.70).

### 3.6 Relationship between variables

#### 3.6.1 IAcc and Singing Experience

A linear regression analysis was conducted to examine whether accumulated hours of singing training predict interoceptive accuracy (IAcc) across amateur and professional singers, with sex included as a covariate. The model excluded age due to its high correlation with total hours of singing, thereby reducing multicollinearity concerns. The overall regression model was significant, F(2,44) = 3.98, p = 0.03, and explained approximately 15% of the variance in IAcc (R² = 0.15, adjusted R² = 0.11). Total hours of singing significantly predicted IAcc (b = 0.00073, SE = 0.00034, t = 2.15, p = 0.04), with higher accumulated hours associated with greater interoceptive accuracy (Figure 3A). Sex also significantly predicted IAcc (b = 8.02, SE = 3.84, t = 2.09, p = 0.04), with males exhibiting higher IAcc compared to females.

**Figure 3:**
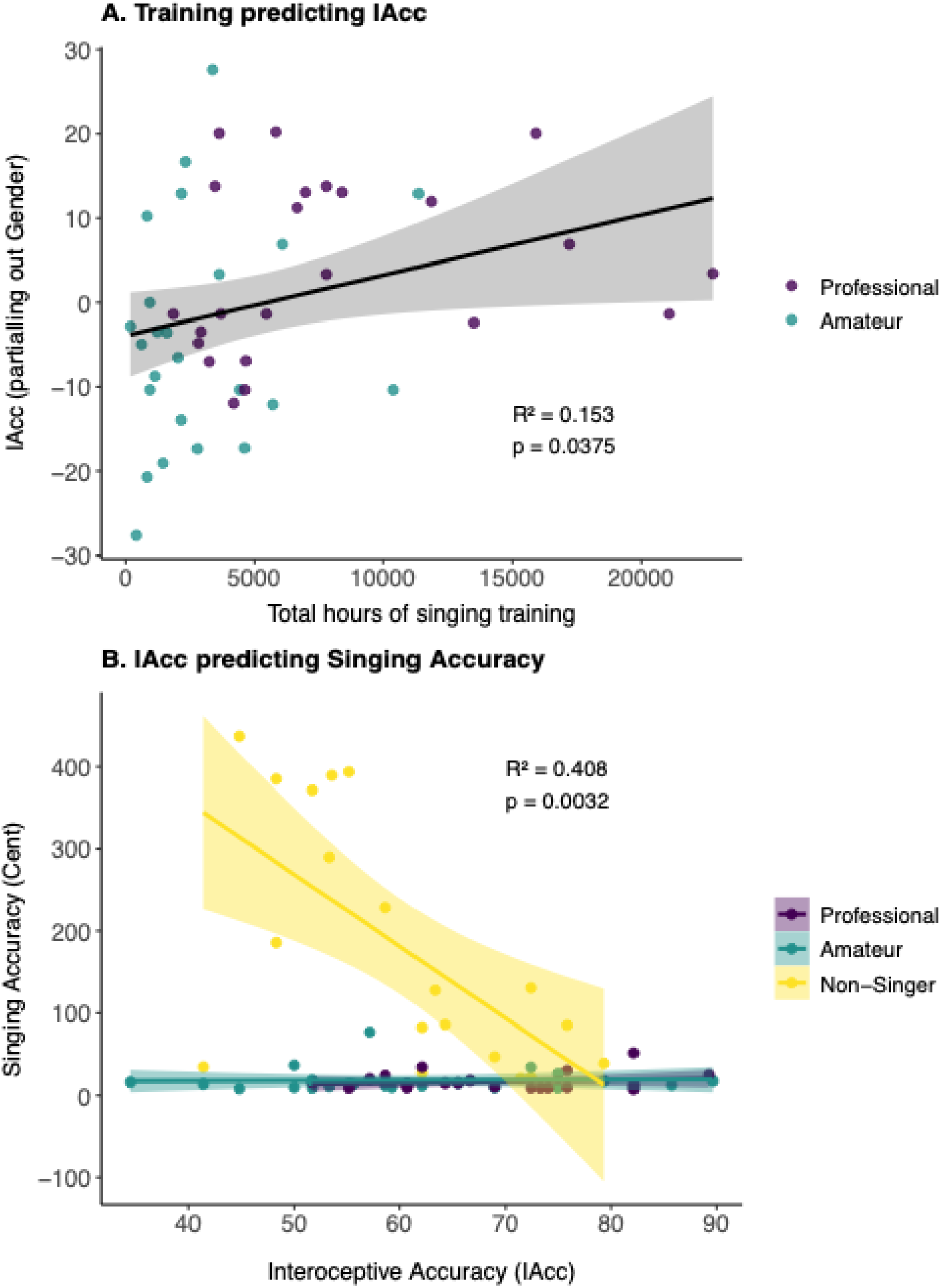
(A) Scatterplot with a linear regression line illustrating the relationship between total accumulated singing hours (Total h/sing) and interoceptive accuracy (IAcc) among professional and amateur singers, residualized for sex. The solid regression line represents the relationship, while the shaded area indicates the 95% confidence interval. A positive trend is observed, with greater accumulated singing hours significantly predicting enhanced interoceptive accuracy (R² = 0.15, p = 0.03). (B) Scatterplot showing the relationship between interoceptive accuracy (IAcc) and pitch accuracy (deviation from target pitch in cents) across non-singers, amateurs, and professionals. Separate regression lines for each group are displayed, with confidence bands. The interaction term between IAcc and singing experience was significant, revealing a strong negative relationship between IAcc and deviation from target pitch in non-singers (R² = 0.41, p = 0.003), but no significant relationship in amateurs (R² = 0.0008, p = 0.90) or professionals (R² = 0.026, p = 0.46).

#### 3.6.2 IAcc and Music Competence Outcomes

*Partial correlations* were computed between IAcc and the musical competence tasks (Pitch Matching Accuracy, Equidistant Beat Discrimination Accuracy, METm, METr, and the GOLD-MSI Singing Scale), controlling for age and sex. Significant negative correlations emerged between *IAcc* and *Singing Accuracy* (r = -0.35, *p* = 0.005) and between *IAcc* and *Equidistant Beat Discrimination Accuracy* (r = -0.32, *p* = 0.01). A significant positive correlation was also found between *IAcc* and the *GOLD Singing Scale* (r = 0.26, *p* = 0.04). No significant associations were observed for the MET tests. These findings suggest that better interoceptive accuracy is associated with improved singing ability, reflected in smaller deviations from target pitch, as well as enhanced beat perception, indicated by smaller deviations from optimal beat equidistance, and higher self-reported singing proficiency, independent of age and sex.

*A multiple regression model* was conducted with pitch-matching accuracy as the dependent variable, given its strong correlation with IAcc and its status as the only significant music production task. The model included IAcc, singing experience (non-singer, amateur, professional), and their interaction term, with age and sex as covariates. The overall model was significant, F(6, 59) = 16.19, *p* < 0.001, R² = 0.62, revealing significant main effects of IAcc (b = -6.52, SE = 1.29, t = -5.04, *p* < 0.001) and singing experience. Both amateurs (b = -421.88, SE = 68.74, t = -6.14, *p* < 0.001) and professionals (b = -695.04, SE = 139.55, t = -4.98, *p* < 0.001) showed significantly lower deviation from target pitch than non-singers. Neither age (b = 0.64, p = 0.69) nor sex (b = 7.54, p = 0.40) significantly predicted deviation from target pitch.

The interaction term between IAcc and singing experience was also significant (b = 4.30, SE = 1.06, t = 4.06, *p* < 0.001), indicating that the relationship between IAcc and deviation from target pitch was moderated by singing experience. To further explore this interaction, separate regression analyses were conducted within each group. In non-singers, a significant negative relationship between IAcc and pitch deviation was observed (b = -8.75, SE = 2.56, t = -3.43, *p* = 0.003, R² = 0.41), suggesting a potential link between interoceptive ability and pitch-matching accuracy in individuals without formal vocal training (see Figure 3B). No significant association was found in amateurs (b = 0.03, SE = 0.22, t = 0.13, *p* = 0.90, R² = 0.0008) or professionals (b = 0.16, SE = 0.22, t = 0.75, *p* = 0.46, R² = 0.026).

#### 3.6.3 IAcc and Emotional awareness (MAIA)

Given the significant associations between Emotional Awareness, singing experience, and interoceptive accuracy observed in preceding analyses, we conducted a mediation analysis to determine whether Emotional Awareness mediates the relationship between singing experience and interoceptive accuracy. The mediation model included singing experience as the predictor, Emotional Awareness as the mediator, and IAcc as the outcome variable. Age and sex were not included as covariates, as previous analyses revealed no significant associations with either interoceptive accuracy or emotional awareness. The analysis used bootstrapping with 5,000 samples to estimate the effects.

The direct effect of singing experience on IAcc was not statistically significant, b = 2.75, SE = 1.82, z = 1.51, *p* = .13, 95% CI [-0.89, 6.25], indicating that singing experience alone does not directly predict interoceptive accuracy. The indirect effect of singing experience on IAcc through emotional awareness approached statistical significance, b = 1.60, SE = 0.89, z = 1.80, *p* = 0.07, 95% CI [0.49, 3.38], suggesting a possible mediating role for emotional awareness. The total effect of singing experience on IAcc was statistically significant, b = 4.35, SE = 1.84, z = 2.37, *p* = .02, 95% CI [1.15, 7.57], indicating that singing experience contributes to interoceptive accuracy when considering both direct and indirect pathways.

Path analysis (Figure 4) revealed that singing experience significantly predicted emotional awareness, b = 0.56, SE = 0.18, z = 3.15, *p* = 0.002, 95% CI [0.20, 0.92], suggesting that greater singing expertise is associated with higher emotional awareness. Furthermore, emotional awareness significantly predicted interoceptive accuracy, b = 2.84, SE = 1.12, z = 2.53, *p* = 0.01, 95% CI [0.01, 0.69], supporting its role as a psychological mechanism linking singing expertise to interoceptive performance.

**Figure 4:**
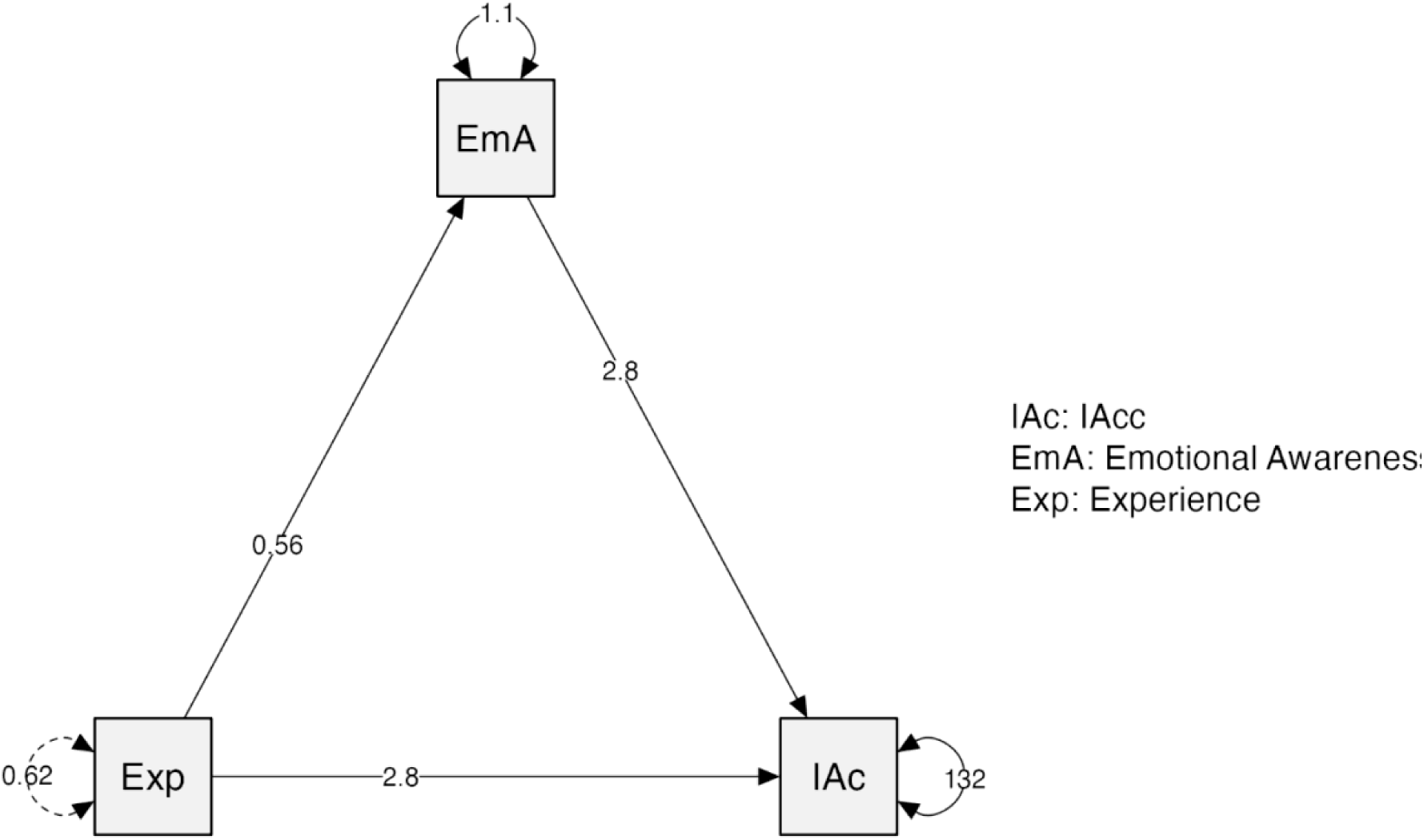
Path analysis examining the mediating role of emotional awareness (EmA) in the relationship between singing experience (Exp) and interoceptive accuracy (IAcc). Solid lines represent explicitly modelled paths, while curved arrows denote variances. Path coefficients are annotated along each arrow. The total effect of singing experience on IAcc was significant, whereas the direct effect remained non-significant when accounting for emotional awareness. The indirect effect through emotional awareness approached significance (*p* = 0.07), suggesting a potential partial mediation.

#### 3.6.4 IAcc and Exteroceptive Temporal Discrimination

A mediation analysis examined whether performance in the exteroceptive control task— assessing auditory-tactile temporal synchrony perception (Control Accuracy; CAcc)— mediates the relationship between singing experience (predictor) and interoceptive accuracy (IAcc; outcome). Age and sex were excluded as covariates, as prior ANCOVA results showed no significant associations with these variables. The analysis used bootstrapping with 5,000 samples to estimate the effects.

The direct effect in the mediation model was not statistically significant, b = 3.19, SE = 1.81, z = 1.76, *p* = .08, 95% CI [-0.33, 6.70]. This suggests that, when accounting for variability in Equidistant Beat Discrimination Accuracy, the unique contribution of singing experience to interoceptive accuracy is reduced. Likewise, the indirect effect through performance in the CAcc task was not significant, b = 1.16, SE = 0.88, z = 1.32, *p* = .19, 95% CI [-0.03, 2.74], indicating that differences in interoceptive accuracy across groups are not driven by differences in temporal estimation abilities. However, the total effect of singing experience on IAcc remained statistically significant, b = 4.35, SE = 1.84, z = 2.37, *p* = 0.02, 95% CI [0.97, 7.43], suggesting that singing experience contributes to interoceptive accuracy through mechanisms other than exteroceptive temporal estimation.

Path analysis (Figure 5) revealed that singing experience significantly predicted performance in the CAcc task, b = 5.41, SE = 1.74, z = 3.10, *p* = 0.002, 95% CI [2.25, 9.18], indicating that participants with greater musical expertise demonstrated superior temporal estimation ability. However, CAcc did not significantly predict IAcc, b = 0.22, SE = 0.16, z = 1.34, *p* = 0.18, 95% CI [-0.03, 0.60], suggesting that while musicians exhibit enhanced temporal judgment, this ability does not explain their interoceptive accuracy.

**Figure 5:**
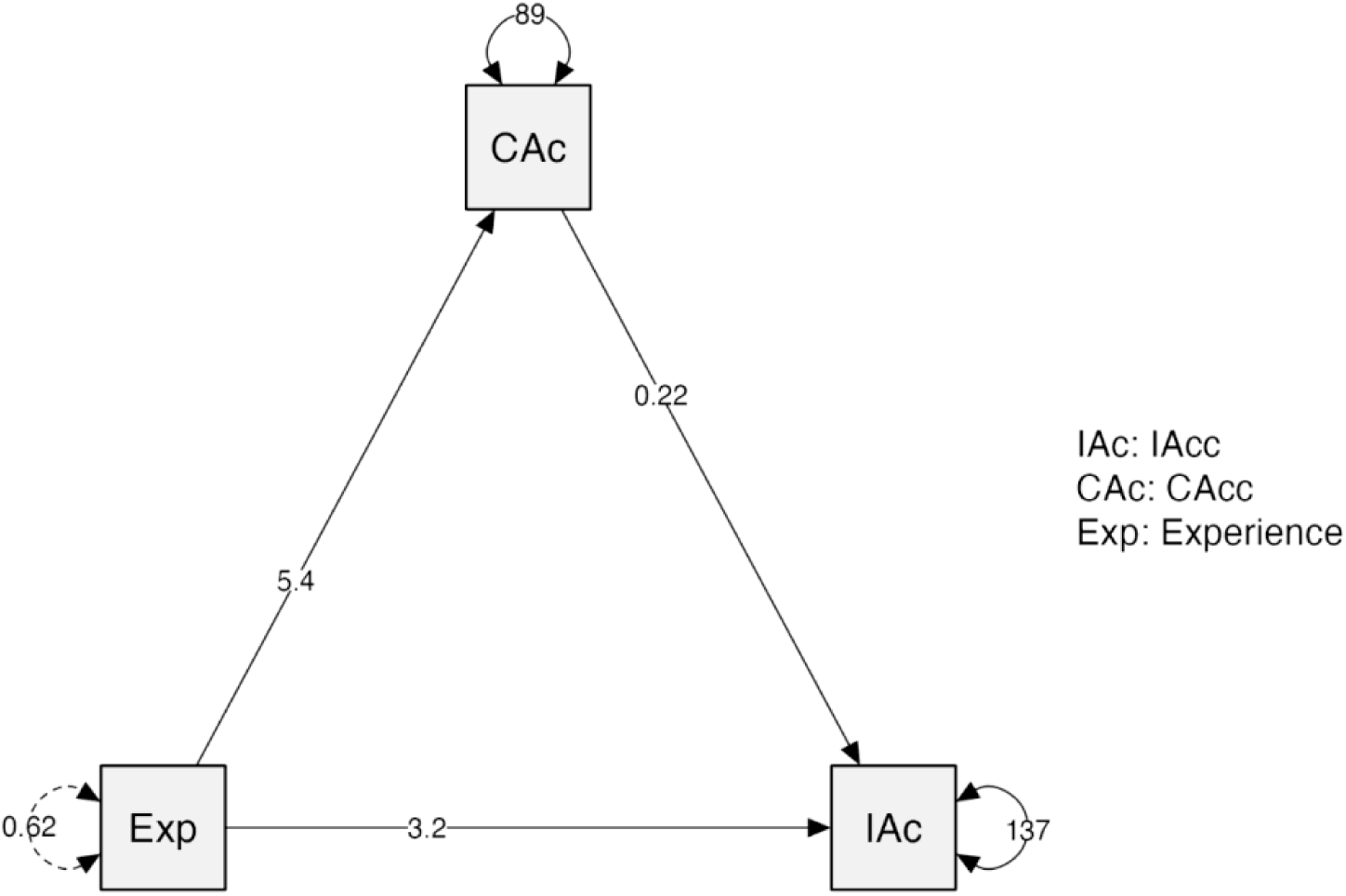
Path analysis examining the mediating role of temporal discrimination ability in the tactile-tone synchrony exteroceptive control task (CAcc) in the relationship between singing experience (Exp) and interoceptive accuracy (IAcc). Solid lines represent explicitly modeled paths, while curved arrows denote variances. Path coefficients are displayed along each arrow. The total effect of singing experience on IAcc was significant, indicating a robust overall association. However, the direct effect became non-significant in the mediation model, suggesting that the unique contribution of singing experience to IAcc is reduced when accounting for CAcc. The indirect effect through CAcc was not significant (*p* = 0.19), indicating that differences in tactile-tonal temporal discrimination ability do not substantially mediate the relationship between singing experience and cardiac IAcc.

#### 3.6.5 IAcc and affective traits

To investigate the relationship between interoceptive accuracy (IAcc) and affective traits, partial correlations were conducted between IAcc, the MINISPIN, and the Trait STAI, controlling for age and sex. The analysis revealed no significant relationship between IAcc and MINISPIN (r=−0.12, *p*=.35) or IAcc and trait STAI (r=0.06, *p*=.65).

## 4. Discussion

In this study, we examined the relationship between interoceptive ability and singing expertise, assessing both objective interoceptive accuracy (IAcc) and self-reported interoceptive sensibility. Our findings suggest that the impact of musical training on IAcc depends on expertise level. Professional singers exhibited significantly higher IAcc than both non-singers and amateur singers, with regression analyses linking accumulated singing hours to greater IAcc. However, this effect was partially mediated by emotional awareness, as assessed through the corresponding MAIA interoceptive sensibility subscale, suggesting that musical training enhances interoception not only through repetitive sensorimotor practice but also through its emphasis on emotional expression.

Beyond training effects, our study provides novel evidence that IAcc relates to musical competence independent of formal training. Non-singers with higher IAcc also demonstrated superior singing accuracy, as indicated by more precise pitch-matching (i.e., smaller deviation from target pitches), suggesting that interoceptive ability may predispose individuals to musical skill acquisition. Altogether, our findings highlight the interplay between innate bodily awareness and experience-dependent plasticity in musical training and help reconcile inconsistencies in prior research by emphasizing expertise over general musical training in interoceptive enhancement.

### 4.1 Singing training as a mechanism to enhance interoception

The role of bodily signals is pronounced in singing, as sound production occurs entirely within the body and depends on precise coordination among multiple motor systems. Volitional breath control, involving the diaphragm and lungs, sustains notes and shapes musical dynamics^43^.

Vocal fold vibration, regulated by air pressure and intrinsic laryngeal muscle activity, determines pitch and sound quality^44^, and timbre, resonance, and intelligibility in lyrical singing are refined through articulatory adjustments of the oral and nasal cavities^45^. Moreover, kinaesthetic awareness and proprioception are crucial for maintaining optimal posture, vocal placement, and pitch control^14,15^. To achieve vocal precision and expressivity, singers must continuously integrate internal bodily signals with multisensory exteroceptive feedback and motor control processes, anticipating and fine-tuning vocal actions while minimizing reactive corrections^16,17,29,46,47^.

Aligning with previous findings that musicians exhibit enhanced interoceptive abilities compared to non-musicians^19,48^, our study confirms that professional singers possess superior interoceptive accuracy relative to non-singers. Critically, however, our results further refine prior observations by demonstrating that significant improvements in interoceptive accuracy appear contingent upon achieving expert-level proficiency rather than merely engaging in musical activities at an amateur level. This is reflected by the absence of significant differences between amateur singers and non-singers. Although musical expertise did enhance auditory-tactile temporal discrimination performance, this heightened temporal sensitivity did not significantly contribute to interoceptive accuracy, thus ruling out a temporal estimation bias as a potential explanation.

Supporting the role of training intensity, the total accumulated singing hours significantly predicted interoceptive accuracy among trained singers, with professionals engaging in nearly three times the annual training hours of amateurs. However, the modest explanatory power, accounting for only approximately 14% of its variance, suggests that additional factors beyond mere training quantity influence interoceptive ability. One possible explanation is the continuous monitoring of bodily feedback, which forms the basis for fine-tuning vocal production during singing training^13,49^. Singers typically begin formal training later than instrumentalists—after voice maturation—limiting the total period available for structured practice^19,50,51^. The heightened reliance on bodily signals may compensate for this shorter training window. However, individual differences in body perception may influence how much singers’ training affects IAcc, with some potentially more attuned to bodily signals from the outset^52^, which could lead to greater gains.

When accounting for other key factors influencing IAcc and singing expertise, emotional awareness emerged as a significant mediator. Greater singing expertise predicted higher emotional awareness, which, in turn, was associated with enhanced IAcc^53,54^. The finding that professional singers exhibited significantly greater emotional awareness than non-singers further supports the role of singing as an embodied practice integrating physiological regulation with affective self-awareness. Indeed, singing—and musical expertise more broadly—requires not only the mastery of fine motor control but also emotional expressivity, both of which are essential for delivering a compelling performance. Expert vocalists must execute complex motor techniques with precision to maintain accurate pitch, timing, and articulation, while simultaneously conveying both the composer’s intended emotions and their own interpretative expression^55^. This dual demand distinguishes high-level performance from mere technical proficiency, as expressivity depends not only on mechanical accuracy but also on the ability to modulate dynamics, phrasing, and timbre in response to emotional cues^56–60^. Moreover, beyond outward emotional expression, musicians must regulate their own emotional states during performance. Performance anxiety, emotional arousal, and cognitive load can interfere with motor control and expressive delivery, making emotional self-regulation a critical aspect of expertise^61,62^. Skilled performers develop coping strategies—such as breath control, attentional focus, and predictive motor planning—to maintain expressive fluidity without compromising technical precision^63^. Our findings underscore this interplay, showing that long-term singing training not only enhances IAcc but also cultivates heightened emotional awareness, which in turn supports interoceptive processing.

Our results contrast with those of Herman et al.^21^, who found no significant differences in interoceptive accuracy, insight, or sensibility between musicians and non-musicians. However, their resting-state fMRI analysis revealed increased sensorimotor network connectivity in musicians, suggesting that musical training may still shape brain regions involved in interoceptive processing. One likely explanation for this discrepancy lies in differences in sample characteristics. While our study included both male and female participants, Herman et al.^21^ exclusively examined female musicians. Although our study did not find significant sex differences in IAcc between groups, the regression analysis revealed that sex independently predicted interoceptive accuracy, indicating that males generally exhibited higher IAcc scores than females when accounting for accumulated singing experience. This aligns with a recent meta-analysis, suggesting that males generally show superior interoceptive accuracy^64^. Additionally, our male participants reported greater confidence in their cardioceptive perception, indicating subtle differences in interoceptive awareness that warrant further investigation.

Another key difference is the type of musical training examined. Unlike keyboard instrumentalists, singers must actively monitor breath control, laryngeal muscle tension, and vocal tract configuration to produce expressive vocalizations—bodily signals closely tied to interoceptive processing. Schirmer-Mokwa et al.^19^ found that professional singers and string players exhibited superior interoceptive accuracy compared to non-musicians. However, accumulated musical practice predicted interoceptive accuracy only in singers, not in string players, possibly due to a ceiling effect. In contrast, Herman et al.^21^ did not find significant group differences in interoceptive accuracy between keyboard players and non-musicians. This discrepancy may reflect differences in the type of musical training, with singers relying more directly on bodily feedback, while keyboard training may provide less interoceptive engagement.

Finally, training onset, intensity, and expertise level differed substantially between studies. Our sample included both professional and amateur singers, whereas Herman et al.^21^ focused on keyboard players with formal secondary music education and a longitudinal cohort exposed to only six months of piano training. While Herman et al.’s musicians had 11–20 years of musical training, they were not professional-level performers, which may explain the absence of interoceptive advantages. In contrast, our sample included professional singers who engage in more intensive and embodied training, likely enhancing their reliance on interoceptive feedback. This aligns with our findings that amateur singers did not exhibit significantly greater IAcc than non-singers, suggesting that moderate training alone may be insufficient to induce measurable interoceptive changes. The difference in training intensity, the level of expertise, and the degree of bodily engagement may furthermore account for the discrepancies between studies.

### 4.2 Interoception accuracy and singing: nurture and nature?

A key question in understanding the relationship between interoceptive accuracy (IAcc) and singing ability is whether heightened interoception facilitates vocal control from the outset or whether extensive singing training actively refines interoceptive precision. Our findings contribute to this debate by showing that while longer accumulated singing training is associated with enhanced IAcc, non-singers with higher interoceptive accuracy also exhibited better pitch-matching abilities. Intriguingly, this suggests that individual differences in bodily awareness may contribute to singing accuracy even in the absence of formal training but does not establish causality.

Interestingly, previous research defined singing accuracy and perceptual pitch discrimination as primary determinants of untrained singing talent, yet also questioned whether pitch perception is sufficient for accurate singing. While some studies have found a correlation between perceptual pitch discrimination and singing accuracy in non-musicians^65–67^, Zarate et al.^68^ demonstrated that pitch-discrimination training alone does not improve singing accuracy. This suggests that perceptual abilities enhance singing ability only when integrated with motor mechanisms, necessitating continuous integration of auditory, somatosensory, and interoceptive feedback^66,69,70^. Given that IAcc reflects an individual’s ability to monitor internal physiological states^1,2^, heightened interoception could facilitate subtle vocal adjustments, enhancing pitch control^71^. However, the exact nature of this relationship requires further exploration.

Trained singers did not show a significant correlation between IAcc and pitch-matching, likely due to a ceiling effect, as their high levels of musical competence reduced inter-individual variability in performance. However, this does not imply that physiological feedback is irrelevant in this group. Kleber et al.^17^ found that professional vocalists can maintain pitch accuracy even when auditory feedback is masked, while Kleber et al.^16^ demonstrated that disrupting physiological input from the larynx adversely affects their pitch accuracy, emphasizing the importance of bodily feedback in vocal motor control. These findings indicate that expert singers rely on well-integrated physiological feedback for pitch regulation, and while predictive control mechanisms may assist in compensating for unreliable sensory input, they do not necessarily replace the role of body awareness in trained vocalists.

Our findings highlight a dynamic interplay between interoceptive ability, musical competence, and vocal training. While extensive singing training enhances interoceptive accuracy, individual differences in bodily awareness may contribute to singing accuracy—measured through pitch-matching—even without formal training, suggesting that pre-existing interoceptive sensitivity provides an initial advantage in vocal control. However, long-term practice remains essential for refining interoceptive precision in expressive singing. Genetic research supports the role of both innate and environmental influences in vocal ability^72,73^, showing that genetic (∼40%) and shared environmental (∼37%) factors contribute equally to singing ability, while early musical experiences—not just formal training—can shape vocal skill development.

## 5. Conclusion

Altogether, this study contributes to a growing body of research on interoception in skilled musical performance, underscoring expertise level as a key determinant and emotional awareness as a mediator, reinforcing interoception’s role in affective and sensorimotor integration. Future studies should examine the neural mechanisms underlying these effects, explore how long-term singing interventions enhance interoceptive function, and track how interoceptive awareness evolves with expertise. Longitudinal research is also needed to determine whether individual differences in interoception predict singing accuracy and how genetic or environmental factors shape this relationship. By advancing our understanding of the physiological factors behind expert musicianship, these insights open new directions for research at the intersection of sensory awareness, emotional regulation, and performance skills.

## Acknowledgements

This study was funded by the Danish National Research Foundation (DNRF117) and the Carlsberg Foundation (CF22-1172).

## Author contribution

BK, PV, EB, and AZ contributed to the conceptualization and design of the study. BK, NT, and AZ were involved in the experiment setup. BK recruited participants and collected the data. BK, AZ, NT and PM contributed to the data analysis. BK and AZ drafted the article. DL contributed to an earlier version of the manuscript and initial analyses. All authors discussed the results, contributed to the interpretation, and commented on the manuscript.

## Notes

### Competing Interest Statement

The authors have declared no competing interest.

## Bibliography

1. Ceunen E., J.W.S. Vlaeyen & I. Van Diest. 2016. On the origin of interoception. Frontiers in Psychology 7: 743. 10.3389/FPSYG.2016.00743/BIBTEX

2. Craig A.D. 2002. How do you feel? Interoception: the sense of the physiological condition of the body. Nature reviews. Neuroscience 3: 655–66. 10.1038/nrn894

3. Berntson G.G. & S.S. Khalsa. 2021. Neural Circuits of Interoception. Trends in Neurosciences 44: 17–28. 10.1016/j.tins.2020.09.011

4. Feldman M.J., E. Bliss-Moreau & K.A. Lindquist. 2024. The neurobiology of interoception and affect. Trends in Cognitive Sciences 28: 643–661. 10.1016/j.tics.2024.01.009

5. Feldman M.J., T.A. Jolink, G.M. Alvarez, et al. 2023. The roles of inflammation, affect, and interoception in predicting social perception. Brain, Behavior, and Immunity 112: 246–253. 10.1016/j.bbi.2023.05.011

6. MacCormack J.K. & K.A. Lindquist. 2019. Feeling hangry? When hunger is conceptualized as emotion. Emotion 19: 301–319. 10.1037/emo0000422

7. Criscuolo A., V. Pando-Naude, L. Bonetti, et al. 2022. An ALE meta-analytic review of musical expertise. Sci Rep 12: 11726. 10.1038/s41598-022-14959-4

8. Furuya S., T. Oku, F. Miyazaki, et al. 2015. Secrets of virtuoso: neuromuscular attributes of motor virtuosity in expert musicians. Scientific Reports 5: 15750. 10.1038/srep15750

9. Zatorre R.J., J.L. Chen & V.B. Penhune. 2007. When the brain plays music: auditory-motor interactions in music perception and production. Nature reviews. Neuroscience 8: 547–558. 10.1038/nrn2152

10. Keough M., D. Derrick & B. Gick. 2019. Cross-Modal Effects in Speech Perception. Annual Review of Linguistics 5: 49–66. 10.1146/annurev-linguistics-011718-012353

11. Kleber B., R. Veit, N. Birbaumer, et al. 2010. The brain of opera singers: experience-dependent changes in functional activation. Cereb Cortex 20: 1144–1152. 10.1093/cercor/bhp177

12. Zamorano A.M., R.J. Zatorre, P. Vuust, et al. 2023. Singing training predicts increased insula connectivity with speech and respiratory sensorimotor areas at rest. Brain Research 1813: 148418. 10.1016/j.brainres.2023.148418

13. Kent R.D. 2024. The Feel of Speech: Multisystem and Polymodal Somatosensation in Speech Production. Journal of Speech, Language, and Hearing Research 67: 1424–1460. 10.1044/2024_JSLHR-23-00575

14. Lindblom B.E. & J. Sundberg. 1971. Acoustical consequences of lip, tongue, jaw, and larynx movement. J Acoust Soc Am 50: 1166–79.

15. Lindblom B.E. & J. Sundberg. 2014. The Human Voice in Speech and Singing. In Springer Handbook of Acoustics Rossing T.D., Ed. 703–746. New York, NY: Springer. 10.1007/978-1-4939-0755-7_16

16. Kleber B., A.G. Zeitouni, A. Friberg, et al. 2013. Experience-dependent modulation of feedback integration during singing: role of the right anterior insula. J Neurosci 33: 6070–80. 10.1523/JNEUROSCI.4418-12.2013

17. Kleber B., A. Friberg, A. Zeitouni, et al. 2017. Experience-dependent modulation of right anterior insula and sensorimotor regions as a function of noise-masked auditory feedback in singers and nonsingers. NeuroImage 147: 97–110. 10.1016/j.neuroimage.2016.11.059

18. Kleber B. & J.M. Zarate. 2014. The Neuroscience of Singing. In The Oxford Handbook of Singing Graham W. & Nix J., Eds. Oxford, UK.: Oxford University Press. 10.1093/oxfordhb/9780199660773.013.015

19. Schirmer-Mokwa K.L., P.R. Fard, A.M. Zamorano, et al. 2015. Evidence for Enhanced Interoceptive Accuracy in Professional Musicians. Front Behav Neurosci 9: 349. 10.3389/fnbeh.2015.00349

20. Hina F., J. Aspell & F. Cardini. 2020. Enhanced behavioural and brain responses to interoceptive signals in musicians.. 10.31234/osf.io/smdwc

21. Herman A.M., A. Olszewska, M. Gaca, et al. 2023. Interoception and the musical brain: Evidence from cross-sectional and longitudinal behavioral and resting-state fMRI study. Psychophysiology 60: e14402. 10.1111/psyp.14402

22. Garfinkel S.N., A.K. Seth, A.B. Barrett, et al. 2015. Knowing your own heart: distinguishing interoceptive accuracy from interoceptive awareness. Biol Psychol 104: 65–74. 10.1016/j.biopsycho.2014.11.004

23. Mehling W.E., M. Acree, A. Stewart, et al. 2018. The Multidimensional Assessment of Interoceptive Awareness, Version 2 (MAIA-2). PLOS ONE 13: e0208034. 10.1371/journal.pone.0208034

24. Faul F., E. Erdfelder, A. Buchner, et al. 2009. Statistical power analyses using G*Power 3.1: tests for correlation and regression analyses. Behav Res Methods 41: 1149–1160. 10.3758/BRM.41.4.1149

25. Christensen J.F., S.B. Gaigg & B. Calvo-Merino. 2017. I can feel my heartbeat: Dancers have increased interoceptive accuracy. Psychophysiology. 10.1111/psyp.13008

26. Brener J. & C. Kluvitse. 1988. Heartbeat detection: judgments of the simultaneity of external stimuli and heartbeats. Psychophysiology 25: 554–561.

27. Kleckner I.R., J.B. Wormwood, W.K. Simmons, et al. 2015. Methodological recommendations for a heartbeat detection-based measure of interoceptive sensitivity. Psychophysiology 52: 1432–1440. 10.1111/psyp.12503

28. Wiens S. & S.N. Palmer. 2001. Quadratic trend analysis and heartbeat detection. Biological Psychology 58: 159–175. 10.1016/S0301-0511(01)00110-7

29. Kleber B., C. Dale, A.M. Zamorano, et al. 2025. Increased Callosal Thickness in Early Trained Opera Singers. 2025.02.28.640273. 10.1101/2025.02.28.640273

30. Mehling W.E., C. Price, J.J. Daubenmier, et al. 2012. The Multidimensional Assessment of Interoceptive Awareness (MAIA). PLOS ONE 7: e48230. 10.1371/journal.pone.0048230

31. Coffey E.B.J., S.C. Herholz, S. Scala, et al. 2011. Montreal Music History Questionnaire: a tool for the assessment of music-related experience in music cognition research. In The Neurosciences and Music IV: Learning and Memory, Conference. Edinburgh, UK.

32. Wallentin M., A.H. Nielsen, M. Friis-Olivarius, et al. 2010. The Musical Ear Test, a new reliable test for measuring musical competence. Learning and Individual Differences 20: 188–196. 10.1016/j.lindif.2010.02.004

33. Levitt H. 1971. Transformed up-down methods in psychoacoustics. J Acoust Soc Am 49: Suppl 2:467+.

34. Wallman-Jones A., P. Perakakis, M. Tsakiris, et al. 2021. Physical activity and interoceptive processing: Theoretical considerations for future research. Int J Psychophysiol 166: 38–49. 10.1016/j.ijpsycho.2021.05.002

35. Wallman-Jones A., E.R. Palser, V. Benzing, et al. 2022. Acute physical-activity related increases in interoceptive ability are not enhanced with simultaneous interoceptive attention. Sci Rep 12: 15054. 10.1038/s41598-022-19235-z

36. Lee P.H., D.J. Macfarlane, T.H. Lam, et al. 2011. Validity of the international physical activity questionnaire short form (IPAQ-SF): A systematic review. Int J Behav Nutr Phys Act 8: 1–11. 10.1186/1479-5868-8-115

37. Domschke K., S. Stevens, B. Pfleiderer, et al. 2010. Interoceptive sensitivity in anxiety and anxiety disorders: an overview and integration of neurobiological findings. Clinical psychology review 30: 1–11. 10.1016/j.cpr.2009.08.008

38. Stevens S., A.L. Gerlach, B. Cludius, et al. 2011. Heartbeat perception in social anxiety before and during speech anticipation. Behaviour research and therapy 49: 138–43. 10.1016/j.brat.2010.11.009

39. Simoens V.L., S. Puttonen & M. Tervaniemi. 2015. Are music performance anxiety and performance boost perceived as extremes of the same continuum? Psychology of Music 43: 171–187.

40. Spielberger C.D. 1989. “State-trait anxiety inventory : a comprehensive bibliography,”, 2nd ed. Palo Alto, CA (577 College Ave., Palo Alto 94306): Consulting Psychologists Press.

41. Connor K.M., K.A. Kobak, L.E. Churchill, et al. 2001. Mini-SPIN: A brief screening assessment for generalized social anxiety disorder. Depress Anxiety 14: 137–40.

42. Allen M., D. Poggiali, K. Whitaker, et al. 2021. Raincloud plots: a multi-platform tool for robust data visualization. Wellcome Open Res 4: 63. 10.12688/wellcomeopenres.15191.2

43. Leanderson R., J. Sundberg & C. von Euler. 1987. Role of diaphragmatic activity during singing: a study of transdiaphragmatic pressures. Journal of applied physiology (Bethesda, Md. 62: 259–270.

44. Shipp T., E.T. Doherty & S. Haglund. 1990. Physiologic factors in vocal vibrato production. Journal of Voice 4: 300–304. 10.1016/S0892-1997(05)80045-1

45. Sundberg J., P. Birch, B. Gumoes, et al. 2007. Experimental findings on the nasal tract resonator in singing. Journal of Voice 21: 127–137.

46. Jones J.A. & D. Keough. 2008. Auditory-motor mapping for pitch control in singers and nonsingers. Exp Brain Res 190: 279–87. 10.1007/s00221-008-1473-y

47. Mürbe D., F. Pabst, G. Hofmann, et al. 2002. Significance of auditory and kinesthetic feedback to singers’ pitch control. J Voice 16: 44–51.

48. Hina F., J. Aspell & F. Cardini. 2020. Enhanced behavioural and brain responses to interoceptive signals in musicians.. 10.31234/osf.io/smdwc

49. Cohen A.J., D.J. Levitin & B. Kleber. 2020. Brain mechanisms underlying singing. In The Routledge Companion to Interdisciplinary Studies in Singing, Volume I: Development Russo F.A., Ilari B., & Cohen A.J., Eds. 79–86. Routledge.

50. Kleber B., N. Birbaumer, R. Veit, et al. 2007. Overt and imagined singing of an Italian aria. NeuroImage 36: 889–900. 10.1016/j.neuroimage.2007.02.053

51. Schlaug G., L. Jancke, Y. Huang, et al. 1995. In vivo evidence of structural brain asymmetry in musicians. Science 267: 699–701.

52. Lametti D.R., S.M. Nasir & D.J. Ostry. 2012. Sensory preference in speech production revealed by simultaneous alteration of auditory and somatosensory feedback. J Neurosci 32: 9351–8. 10.1523/JNEUROSCI.0404-12.2012

53. Critchley H.D. & S.N. Garfinkel. 2017. Interoception and emotion. Current Opinion in Psychology 17: 7–14. 10.1016/j.copsyc.2017.04.020

54. Schuette S.A., N.L. Zucker & M.J. Smoski. 2021. Do interoceptive accuracy and interoceptive sensibility predict emotion regulation? Psychological Research 85: 1894–1908. 10.1007/s00426-020-01369-2

55. Coutinho E., K.R. Scherer & N. Dibben. 2019. Singing and emotion. In The Oxford handbook of singing Welch G., Howard D., & Nix J., Eds. 296–314. Oxford, UK: Oxford University Press.

56. Juslin P.N. & P. Laukka. 2003. Emotional expression in speech and music: evidence of cross-modal similarities. Ann N Y Acad Sci 1000: 279–82.

57. Scherer K.R. 1995. Expression of emotion in voice and music. J Voice 9: 235–48.

58. Scherer K.R. 2013. The singer’s paradox: on authenticity in emotional expression on the opera stage. In The Emotional Power of Music Cochrane T., Fantini B., & Scherer K.R., Eds. 55–73. Oxford: Oxford University Press.

59. Scherer K.R., T. Johnstone & G. Klasmeyer. 2003. Vocal expression of emotion. In Handbook of the Affective Sciences Davidson R.J., Scherer K.R., & Goldsmith. H, Eds. 433–456. New York: Oxford University Press,.

60. Scherer K.R., J. Sundberg, B. Fantini, et al. 2017. The expression of emotion in the singing voice: Acoustic patterns in vocal performance. J Acoust Soc Am 142: 1805. 10.1121/1.5002886

61. Kenny D.T. 2004. Music performance anxiety: is it the music, the performance or the anxiety. 10: 38--43.

62. Thomas J.P. & T. Nettelbeck. 2014. Performance anxiety in adolescent musicians. Psychology of Music 42: 624–634. 10.1177/0305735613485151

63. Fehm L. & K. Schmidt. 2006. Performance anxiety in gifted adolescent musicians. J Anxiety Disord 20: 98–109. 10.1016/j.janxdis.2004.11.011

64. Prentice F. & J. Murphy. 2022. Sex differences in interoceptive accuracy: A meta-analysis. Neurosci Biobehav Rev 132: 497–518. 10.1016/j.neubiorev.2021.11.030

65. Watts C., K. Barnes-Burroughs, M. Andrianopoulos, et al. 2003. Potential factors related to untrained singing talent: a survey of singing pedagogues. Journal of Voice 17: 298–307. 10.1067/S0892-1997(03)00068-7

66. Watts C., R. Moore & K. McCaghren. 2005. The relationship between vocal pitch-matching skills and pitch discrimination skills in untrained accurate and inaccurate singers. Journal of voice : official journal of the Voice Foundation 19: 534–543. 10.1016/j.jvoice.2004.09.001

67. Watts C., J. Murphy & K. Barnes-Burroughs. 2003. Pitch matching accuracy of trained singers, untrained subjects with talented singing voices, and untrained subjects with nontalented singing voices in conditions of varying feedback. Journal of voice : official journal of the Voice Foundation 17: 185–194.

68. Zarate J.M., K. Delhommeau, S. Wood, et al. 2010. Vocal accuracy and neural plasticity following micromelody-discrimination training. PLoS ONE 5: e11181. 10.1371/journal.pone.0011181

69. Moore R.E., J.M. Estis, F. Zhang, et al. 2007. Relations of pitch matching, pitch discrimination, and otoacoustic emission suppression in individuals not formally trained as musicians. Perceptual and motor skills 104: 777–784.

70. Moore R.E., C. Keaton & C. Watts. 2007. The Role of Pitch Memory in Pitch Discrimination and Pitch Matching. Journal of Voice 21: 560–567. 10.1016/j.jvoice.2006.04.004

71. Scotto Di Carlo N. 1994. Internal voice sensitivities in opera singers. Folia Phoniatr Logop 46: 79–85. 10.1159/000266296

72. Yeom D., Y.T. Tan, N. Haslam, et al. 2022. Genetic factors and shared environment contribute equally to objective singing ability. iScience 25: 104360. 10.1016/j.isci.2022.104360

73. Yeom D., N. Haslam, Y.T. Tan, et al. 2024. Twin Data Support a Sensitive Period for Singing Ability. Twin Research and Human Genetics 1–11. 10.1017/thg.2024.30

